# Longitudinal, Multiparametric MRI Assessment of repetitive mild TBI in rats

**DOI:** 10.1101/666024

**Authors:** Ahmad Raza Khan, Brian Hansen, Nina K. Iversen, Jonas L. Olesen, Mariana Angoa-Perez, Donald M. Kuhn, Leif Østergaard, Sune N. Jespersen

## Abstract

Repetitive mild traumatic brain injury (mTBI) has long term health effects and may result in the development of neurodegenerative or neuropsychiatric disorders. Histology shows axonal and dendritic beading, synaptic atrophy, vasodilation and gliosis occuring within hours/days post-mTBI. However, current neuroimaging techniques are unable to detect the early effects of repetitive mTBI. Consequently, mTBI brain scans are normal appearing and inconclusive. Hence, neuroimaging markers capable of detecting subtle microstructural and functional alterations are needed. We present results from longitudinal, multiparametric magnetic resonance imaging (MRI) assessment of repetitive mTBI in rats. We employ advanced in-vivo diffusion MRI (dMRI) to probe brain microstructural alterations, perfusion MRI to assess cerebral blood flow (CBF), close to the injury site, and proton MR spectroscopy to assess metabolic alterations in the ipsilateral cerebral cortex. High resolution anatomical scans were also acquired. In agreement with clinical observations, anatomical scans of rats were normal appearing even after repeated mTBI. Throughout, significance is regarded as p<0.05 post false discovery rate correction. dMRI revealed significant microstructural remodelling in ipsilateral hippocampus (reduced radial kurtosis), may be due to axonal/dendritic beading, demyelination, synaptic atrophy and edema. Consistent with prior reports of reduced cell/fiber density in mTBI, we find significantly increased mean diffusivity in ipsilateral corpus callosum. We also find significantly decreased glutathione (GSH) and increased total Choline (tCho) following second and third mTBI (vs baseline), also reported in clinical mTBI cohorts. Reduced GSH suggests oxidative stress and increase in tCho indicate cell damage/repair. CBF did not change significantly, however, high variability in CBF following the second and third mTBI suggest increased variability in CBF likely due to tissue hypoxia and oxidative stress. Oxidative stress may affect capillary blood flow by disturbing pericyte capillary contraction. Around 40% of pericytes retract after mTBI causing pericyte depletion and white matter dysfunction as suggested by dMRI findings. Multiparametric MRI detects meaningful mTBI-induced alterations otherwise undetectable with conventional MRI. Similar strategies may provide useful information to aid diagnosis of human mTBI.

## 1.0 INTRODUCTION

The incidence of repetitive mild traumatic brain injury (mTBI) is increasing and has been described as an epidemic warranting closer attention [1, 2]. Repetitive mTBI studies supply mounting evidence that these mild injuries have long-term health effects and may result in the development of neurodegenerative diseases [3, 4], depression [5], cognitive impairment [3, 6], personality disorder [7, 8], and accelerated aging [2]. Collectively, mTBI causes disabling ailments and put a large economic burden on society [4, 9]. Repetitive mTBI induces very mild and diffuse a change in damage, which typically cannot be detected by conventional brain scanning methods. However, microscopy has revealed axonal injury [1, 10], dendritic beading and retraction [11, 12], reduction in arteriolar and capillary diameter, and astroglial swelling after mTBI [13–15]. While such immunohistological information is vital to understand the underlying mechanisms of injury, it can only be obtained from animal models and/or cadaver brains. Thus, there is an urgent need for non-invasive neuroimaging probes to detect subtle metabolic, microstructural and pathophysiological alterations for early diagnosis and evaluation of treatment intervention.

Although the early effect of repeated mTBI is not well understood, accumulating evidence prompted many organizations to implement “three-strike” rules, those who suffer a third mTBI should withdraw from the injury surroundings [16]. In the present study we employed a repetitive mTBI (three mTBI) rat model, which imitate human concussion and reflect similar cellular, molecular, and behavioral characteristics of repetitive mTBIs in humans at acute [4, 8], and chronic stages after repetitive mTBI. Multiparametric MRI protocols have the potential to provide markers of subtle tissue injury, metabolic and physiological alterations, which might go undetected if only one of the methods were employed. In the present study, we therefore employed a longitudinal multiparametric MRI approach to look for diffusion MRI (dMRI) based microstructural alteration, ^1^H MRS based metabolic alterations and, perfusion MRI based CBF alteration after repetitive mTBI. Additionally, sucrose consumption test (SCT), carried out weekly before and after repetitive mTBI in rats suggest depressive behavior.

dMRI is a well-established MRI method to probe microstructural alterations. Recently, diffusion kurtosis imaging (DKI) [17], which accounts for non-Gaussian effects in the diffusion signal, has emerged as a clinically feasible method to detect microstructural alterations. DKI has been demonstrated to be more sensitive to microstructural alterations in preclinical disease models [18–21], as well as in mTBI patients [22] compared to conventional diffusion tensor imaging (DTI) parameters. Where conventional DKI (as employed here) uses single diffusion encoding (SDE), another dMRI based method double diffusion encoding (DDE) probes instead spin displacement correlations [23–25]. One DDE parameter, fractional eccentricity (FE) [26, 27] or μFA [28], disentangles the fiber dispersion effects of fractional anisotropy (FA). Another DDE parameter, the eccentricity (ε), is able to probe the shape of the cellular ensemble [26]. Although DTI, DKI and DDE metrics are able to detect subtle microstructural changes, they lack the ability to provide cellular-specificity. Therefore, we also employed ^1^H magnetic resonance spectroscopy (^1^H MRS) in the present study, enabling us to probe cell-specific metabolites such as N-acetyl aspartate (NAA), a neuronal marker [29]; myo-inositol (mI), a glial marker [30]; choline (Cho), a more general markers related to the cell membrane, and the levels of neurotransmitters, γ-aminobutyric acid (GABA), Glutamine (Gln), and Glutamate (Glu). A significant correlation of these metabolic markers with microstructural parameters would complement and enhance our understanding of metabolite and microstructure alterations [31–33]. In addition to microstructural and metabolite perturbations, we also acquired arterial spin labelling (ASL) based perfusion MRI to provide estimates of pathophysiologically relevant parameter, cerebral blood flow (CBF). Other perfusion MRI parameters, cerebral blood volume (CBV) and capillary transit time heterogeneity (CTH) [34] are computable only by perfusion scans with contrast agent. However, these parameters have been found to be altered at preclinical stages of multiple diseases [35–37] and also considered early markers of concussion [38]. Decreased CBF and elevated CTH after repetitive mTBI is believed to provide sufficient blood-tissue oxygen concentration gradient and oxygen exchange time for neuronal function and survival. These changes have been ascribed to compressed capillaries, mitochondrial dysfunction, and glial activation, to name a few [34, 37]. Therefore, perfusion parameters are also crucial to understand early diagnosis and treatment assessment after repetitive mTBI. Such metabolic, microstructural, and CBF derangement of the tissue after repetitive mTBI may contribute to tissue-hypoxia, inflammation, and neurodegeneration, which may further augment secondary damage to the tissue.

The present study revealed significant alterations in the mean diffusivity (MD) estimate from DDE after the third mTBI, and a significant decrease in the radial kurtosis tensor (*W*_T_) from SDE DKI. Our ^1^H MRS experiments showed significant decrease in antioxidant, glutathione level (GSH) and a significant increase in total choline (tCho). Glutamate (Glu) and γ-aminobutyric acid (GABA) levels also increased after mTBI in comparison to baseline. Similar to human concussion, the present study found a marked decrease in CBF in ipsilateral gray matter (injury site) (IGM) of the brain. The reduced AD, increased MD in white matter, increased Glu and lactate are considered potential biomarkers for repetitive/ mTBI, [11, 39, 40]. The MRI based microstructural, metabolic and blood perfusion alterations are corroborated by immunohistological findings in repetitive mTBI studies in animals [41] and humans [42]. By employing DKI, DDE and perfusion MRI in our study, the methods are allowed to support each other and provide complementary information which can potentially be used as objective markers for early diagnosis of repetitive mTBI and for evaluation of new treatment and intervention strategies.

## 2.0 Material and Methods

Male rats (Fisher 344 (Charles River, Germany), n=5, age =10 weeks) were employed after ethical approval by the Danish Animal Ethical Inspectorate, Denmark (2016-15-0201-00877). Animals were acclimatized for two weeks and sucrose consumption test (SCT) were carried out weekly as described previously [43], at baseline and after repetitive mTBI. The repetitive mTBI protocol was adopted from a mouse model developed by Kane et al. 2012 [4] and modified to suit adult rats. The impactor tied with free fishing line (2.2 Kg test, 0.53 mm diameter) was barricaded 1 m above the impact site. Prior to injury, hair on skull were shaved after a brief anesthesia of isoflurane and rat was placed on the horizontal French window platform, of mTBI apparatus. The impactor was released through the perforated PVC guide tube (20 mm diameter and 1.2 m length) and hit the target region of the head. Rat was fallen down on a sponge cushion below the platform after 180° rotation. (Supplementary video 1). The mTBI was performed on the same animals at day1 (TBI1), day3 (TBI2) and day7 (TBI3). Rats were allowed to regain consciousness after impact and then scanned immediately after each mTBI under Isoflurane gas anesthesia. Similar procedure was applied, except mTBI at baseline for MRI scans. MRI scans were performed on anesthetized rat using a 9.4 T MRI system (Bruker, Biospin, Germany) with a BGA-12HP gradient set and a crosscoil combination using a 76mm quadrature coil for excitation and a four element rat brain cryo-surface coil for signal reception (Bruker Biospin, Ettlingen, Germany) as described previously [19, 44].

### 2.1 Diffusion MRI

#### 2.1.1 DKI data

DKI data acquisition was performed as described previously [19]. Briefly, data were acquired according to the fast DKI scheme proposed recently [45–47] with a segmented EPI sequence (two segments), diffusion duration (δ) =4 ms and diffusion separation time (Δ) =12 ms, three b0 images, two b-values 1.0 ms/μm^2^ and 2.5 ms/μm^2^ with nine directions. Other DKI scan parameters were TR/TE 2200/25.8 ms, number of slices =38, isotropic resolution =300 μm, matrix size 128×64, averages=36, resulting in a total scan time of 55 min.

#### 2.1.2 DDE data

DDE data were acquired in the axial-plane using double-pulsed field gradient diffusion experiment scheme as described previously [26] with identical encodings using the following parameters: Δ =10 ms, δ=3 ms, five b0 images, three b-values 0.5, 1, and 2 ms/μm^2^, with 72 directions, TR/TE = 2000/ 65.9 ms, slice thickness=1.2 mm and, in-plane resolution = 0.29 mm. Diffusion-weighted images were acquired separately for each b-value in a scan time of 10 min.

Saturation slices were placed in all orthogonal imaging-planes to reduce the presence of motion artefacts in the dMRI data. Diffusion images were registered to the respective b0 images and were further corrected for noise using the Marchenko-Pastur PCA method [48]. Gibbs ringing artifacts were also corrected by subvoxel shifting [49]. Both DKI and DDE datasets were visually inspected before estimation of DKI and DDE metrics. The anatomical scans were acquired with a rapid acquisition with relaxation enhancement (RARE) sequence, with identical slice position to those of the DKI data and in-plane resolution = 75 μm, matrix size =512×256, TR/TE = 4752/37 ms with two averages and a total scan time of 10 min 8 sec.

#### 2.1.3 Parameter Estimation and ROI placement

Nonlinear least squares fitting was carried out with the Levenberg–Marquardt algorithm implemented in Matlab (The Mathworks, Natick, MA) on pre-processed DKI data. A range of diffusion metrics from kurtosis tensor parameters (MKT, *W*_L_, and *W*_T_) [46, 47, 50] and DTI parameters (AD, RD. MD, and FA) [51, 52] was considered. Post-processed DDE data was also used to compute DTI metrics (AD, RD, MD and FA) and DDE metrics, such as eccentricity (ε), fractional eccentricity (FE) and microscopic anisotropy (MA) [26, 53]. The ROIs were placed manually on the generated FA maps from DKI and DDE dataset on the ipsilateral (I) and contralateral side (C) of the brain respectively, viz. motor cortex (IM and CM), corpus callosum (CCC, ICC), hippocampus (IHP, CHP), caudate putamen (ICP, CCP) and the internal capsule (IIC, CIC) (Figure 1).

**Figure 1:**
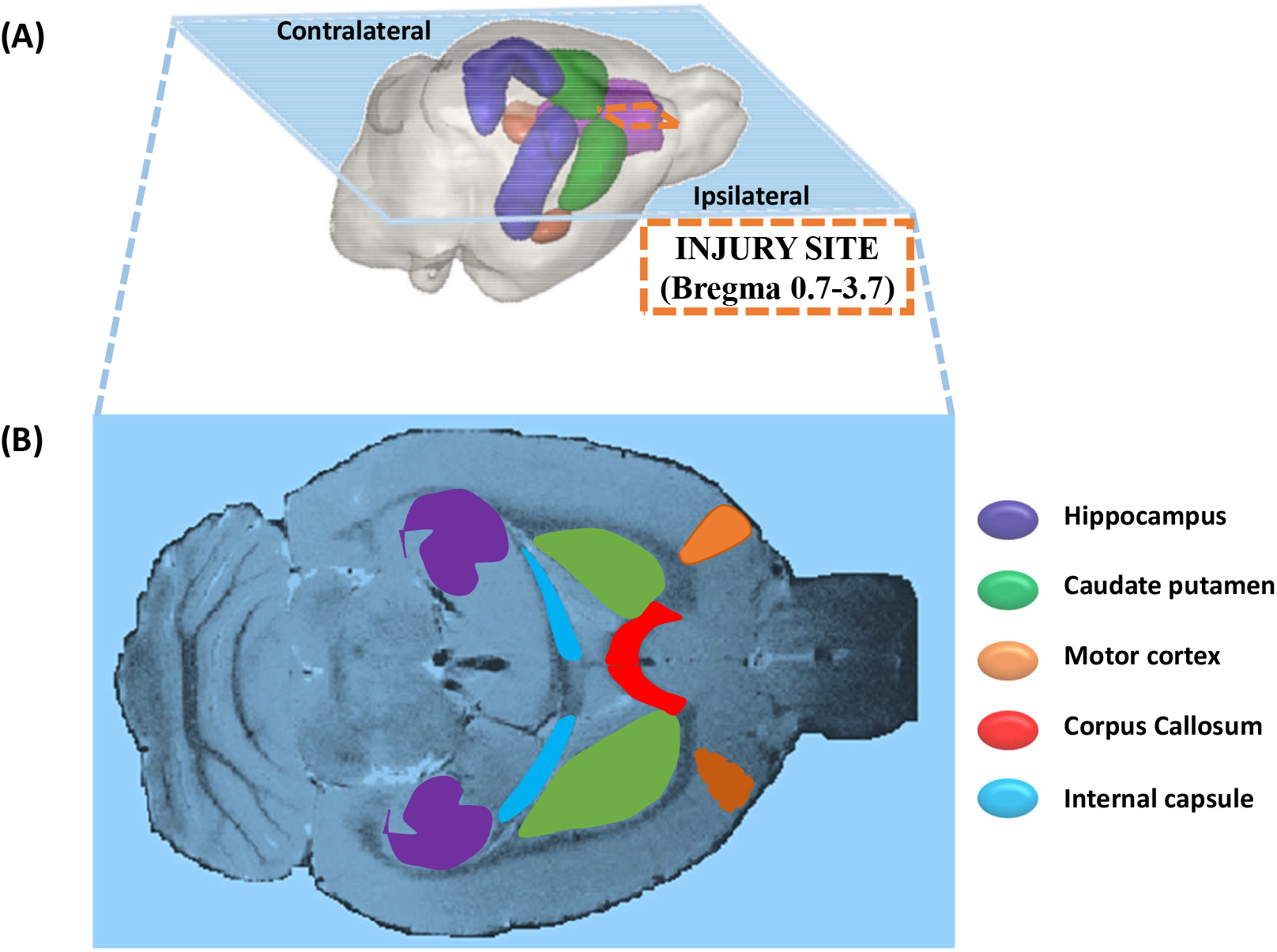
A) 3D representation of a rat brain showing the ipsilateral and contralateral sides of the brain with injury site marked (dashed line). B) A horizontal anatomical image of a rat brain showing targeted ROIs in ipsilateral and contralateral region.

### 2.2 ^1^H MRS

In-vivo ^1^H MRS data were collected from the ipsilateral cerebral cortex of the brain using the point resolved spatially localized spectroscopy (PRESS) sequence with TR/TE 5000/16.5 ms, spectral width 4401 Hz, 256 averages spanning 21 min 20 sec as described previously [19]. Briefly, T2-weighted images were used as reference images to position the volume of interest (Figure 2). Prior to the MRS acquisition, region of interest based shimming (MAPSHIM; Bruker BioSpin, Ettlingen) Spectra were acquired after achieving <20 Hz water signal linewidth (~0.05 ppm) within the voxel and variable power RF pulses with optimized relaxation delays (VAPOR) and outer volume saturation (OVS) scheme were applied to suppress the water signal from the voxel and signal from outside the voxel respectively.

**Figure 2:**
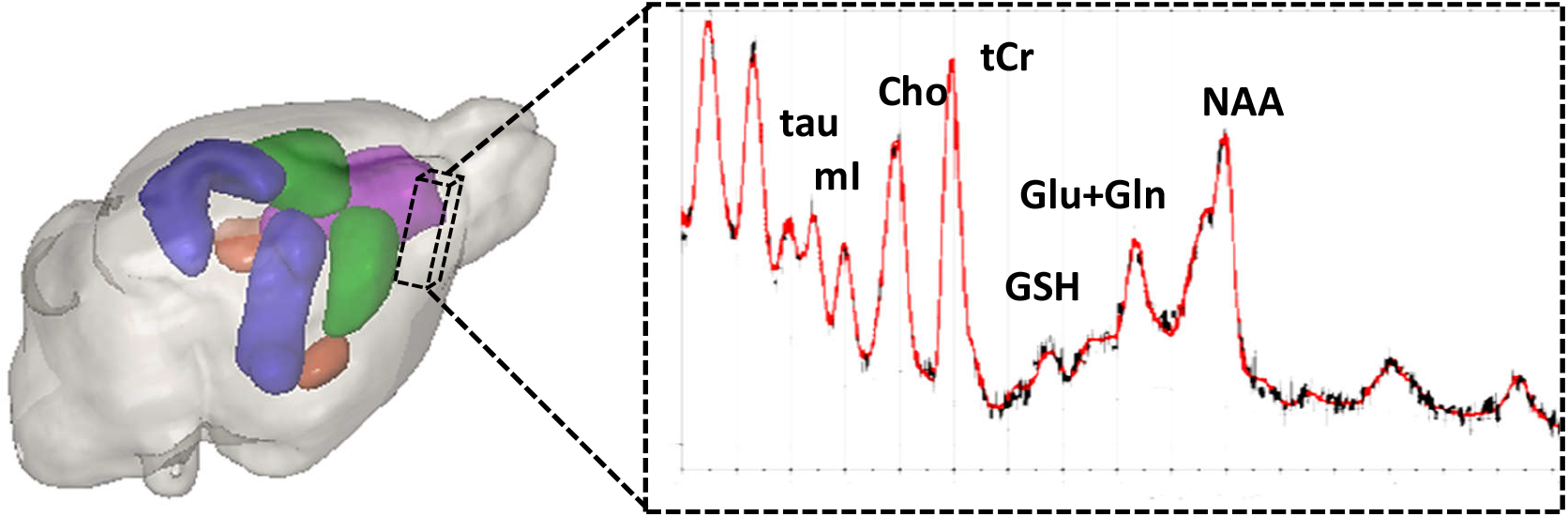
The volume of interest (dashed voxel) 3.5 x1.8 x 4.5 mm ^3^ in the ipsilateral cerebral cortex and a representative ^1^H MR spectrum from the voxel processed in LC Model.

#### 2.2.1 MRS Post-processing using LC Model

Raw ^1^H MRS data were imported into LC model and zero and first-order phase corrections were applied [54]. Quantitative metabolic information was extracted by referencing a basis set obtained from in vitro MR spectra at identical TE and field strength. Further analysis in LC Model provides absolute metabolite concentration and metabolic ratio with respect to total Creatinine (tCr) concentration along with uncertainties of the metabolite concentrations as Cramer-Rao lower bounds (CRLB). These values were used to establish the accuracy of the obtained metabolite concentration. In the present study metabolites having only CRLB <20% are included for further analysis, namely NAA+NAAG, GABA, tCho, mI, Tau, Glu, Gln, and GSH as described previously [19]. Metabolic concentration ratios with respect to tCr were then exported to Matlab for further statistical analysis.

### 2.3 Perfusion MRI

Perfusion MRI data were acquired in the axial-plane using a flow-sensitive alternating recovery Echo Planar Imaging (FAIR-EPI) spin-echo sequence with 16 slice selective (flow dependent) and 16 non-slice selective (flow independent) images using the following parameters: Inversion time (TI) = 100-1600 ms, with 100 ms time interval; TR ≈12 sec; field of view = 32.9×19.2 mm; matrix size = 164×96; 15 slices with 1mm slice thickness.

#### 2.3.1 Parameter Estimation and ROIs placement

Before parameter estimation, denoising [48] and Gaussian smoothing were performed to the inversion recovery (IR) data. T1 maps were generated in Matlab by voxel-wise, non-linear least square fitting (Levenberg-Marquardt) of the IR signal equation to the selective and non-selective IR data thus obtaining T1 maps from both preparations. CBF maps were generated in a voxel-wise manner using measured T1, and a blood tissue partition coefficient of 0.89 ml/gm as described elsewhere [55]. The ROIs were placed manually only on the mid-slice of T1 maps from the perfusion MRI dataset. The ROIs were placed on ipsilateral cortical gray matter (IGM) and white matter (IWM) and also on contralateral gray matter (CGM) and white matter (CWM) region. Only the central slice has well-defined TR in this sequence implementation, therefore we analyzed only the central slice in our analysis. In all cases, the central slice position was intended to coincide with bregma, but some variation was observed during analysis (position span ±1.6mm relative to bregma).

### 2.4 Statistical analysis

The statistical analysis of the multiparametric MRI data was performed using a linear mixed model implemented in Matlab. A voxel-wise analysis was performed for dMRI parameters and CBF parameter and selected metabolites from ^1^H MRS data. Animals were considered as a random effect and group considered as a fixed effect. Level of statistical significance was assessed using an F-test followed with FDR correction as described previously [20].

## 3.0 Results

There were no significant alterations or marked variation in the CP region. Therefore, CP will not be shown in the following results although it was subjected to the same analysis as the other ROIs. Likewise, SDE-DTI metrics (AD, RD, MD, and FA) (Supplementary figure 1 (A-D)) obtained from the DKI analysis as well as DDE metrics (ε, FE and MA) (Supplementary figure 2 (A-C)) did not shown significant alterations in any region at any of the three-time points in comparison to the baseline control. The anatomical images after each mTBI did not show any conspicuous injury to the brain tissue in the ipsilateral cortical region of the brain, although larger ventricles were observed after repetitive mTBI (Figure 3). Sucrose consumption test also shown significant reduction in sucrose consumptions in repetitive mTBI group in comparison to baseline (**p<0.01) (Supplementary figure 3).

**Figure 3:**
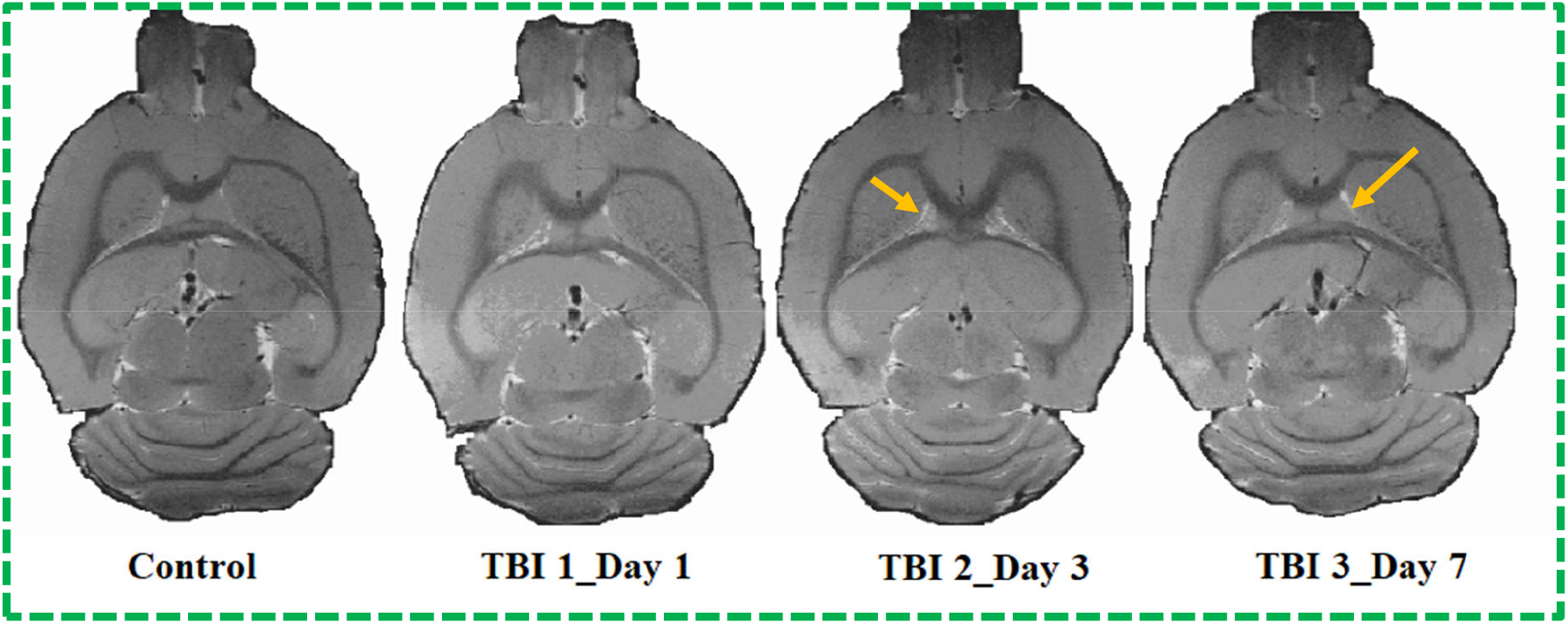
Horizontal T2 weighted-images from a rat brain at baseline (control), after 1^st^ (TBI1_Day1), 2^nd^ (TBI2_Day3) and, 3^rd^ mTBI (TBI3_Day7). Arrows indicate larger ventricle size after repetitive mTBI and no apparent brain injury shown in the ipsilateral cortex.

### 3.1 Diffusion MRI

#### 3.1.1 DKI Parameters (MKT, *W*_L_, *W*_T_)

MKT and *W*_L_ did not show significant alteration even before FDR correction. Only *W*_T_ decreased significantly in the IHP region of the brain after the third mTBI (FDR corrected **p<0.01) (Figure 4). No other alterations were observed in any other ROI in the ipsilateral or contralateral region of the brain. MKT and *W*_L_ metrics did not show marked variation in any region at any three-time points of the study (Figure 5–6).

**Figure 4:**
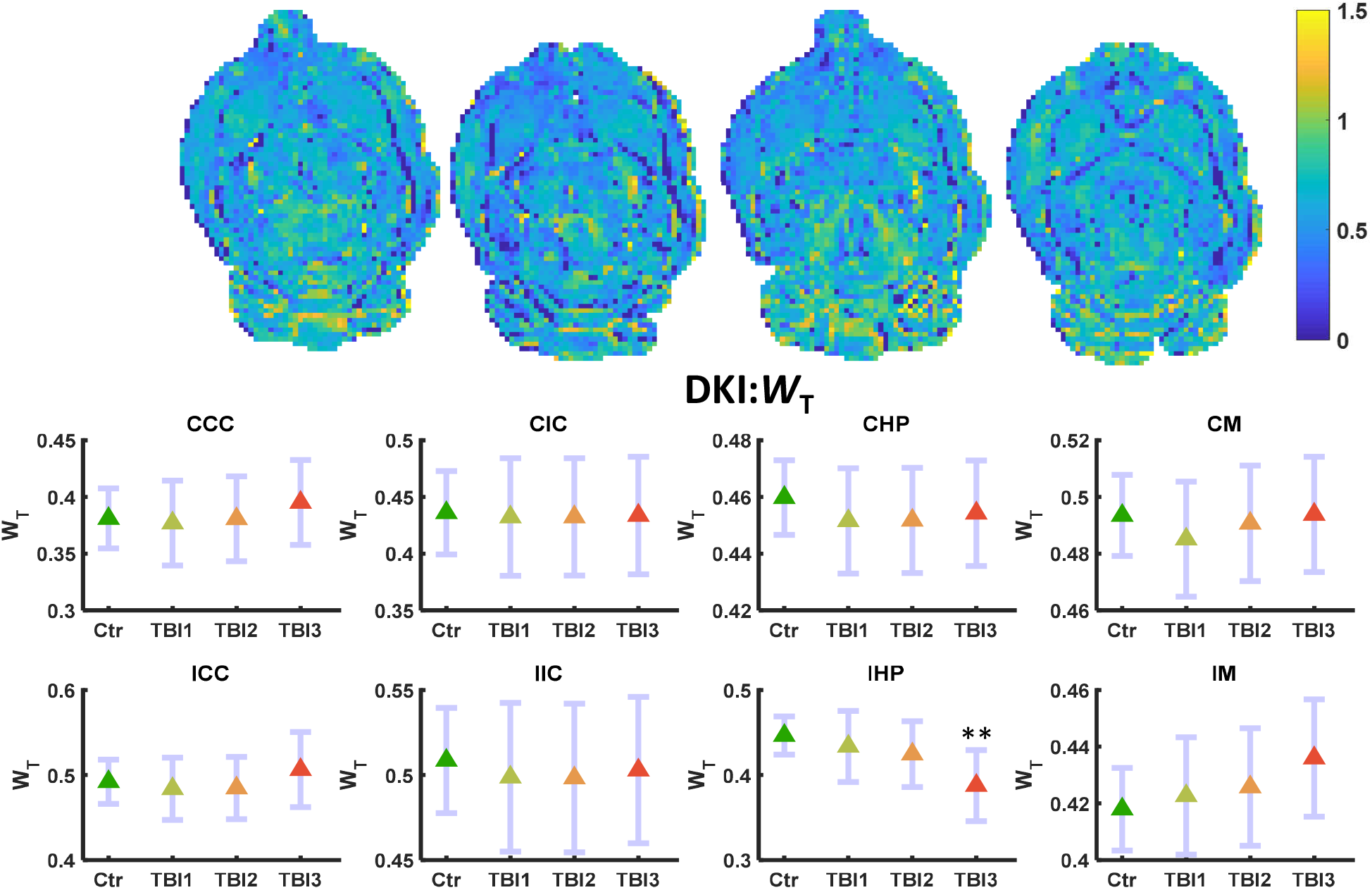
*W*_T_ maps (L-R) control, TBI1, TBI2, and TBI3. Mean (middle) and 95% confidence interval (CI) as error bar in contralateral and ipsilateral ROIs. *W*_T_ decreased significantly (FDR corrected **p<0.01) only in the IHP region of the brain.

**Figure 5:**
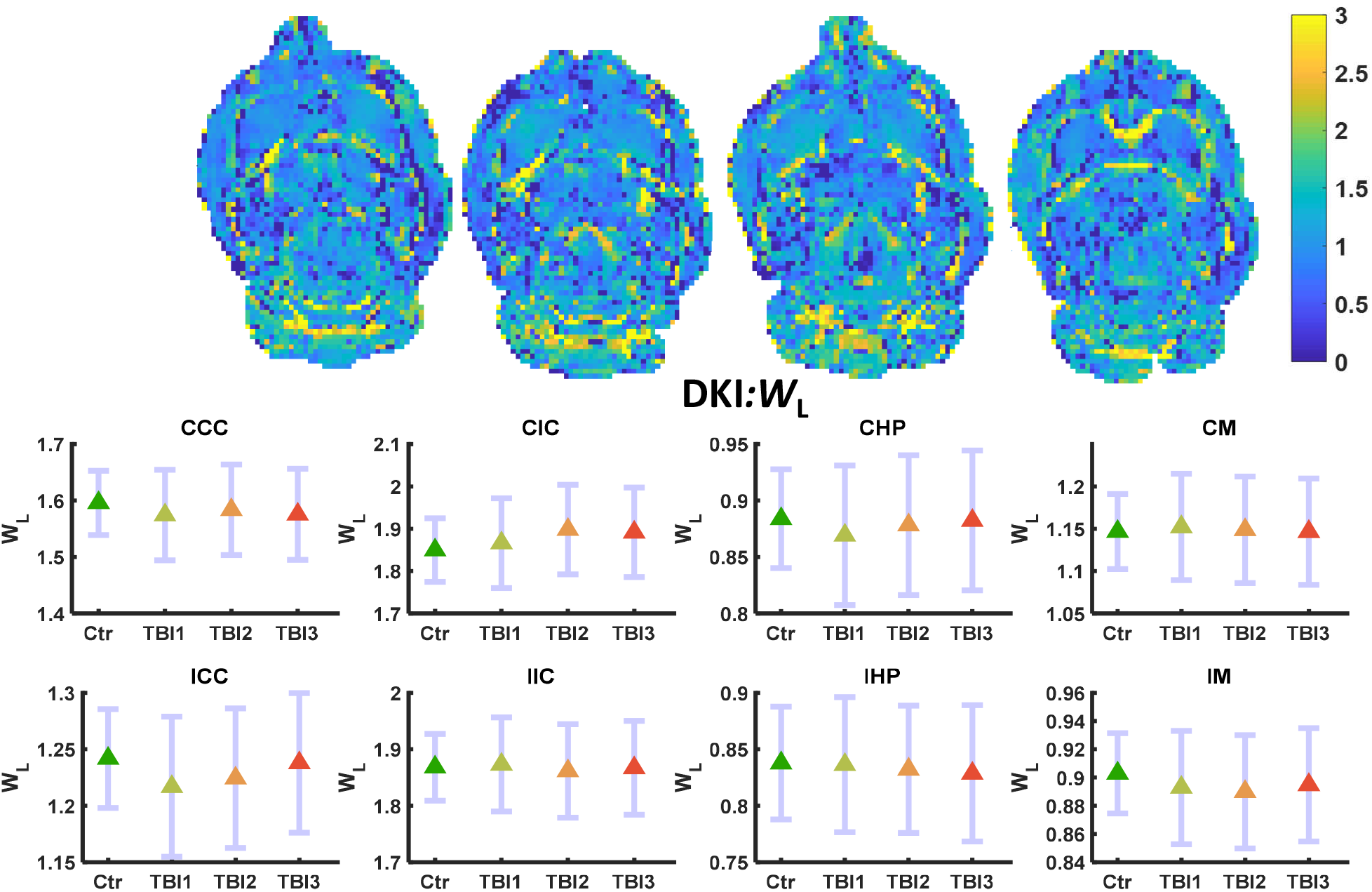
*W*_L_ maps (L-R) control, TBI1, TBI2, and TBI3. Mean (middle) and 95% confidence interval (CI) as error bar in contralateral and ipsilateral ROIs. *W*_L_ did not change significantly in any region of the brain.

**Figure 6:**
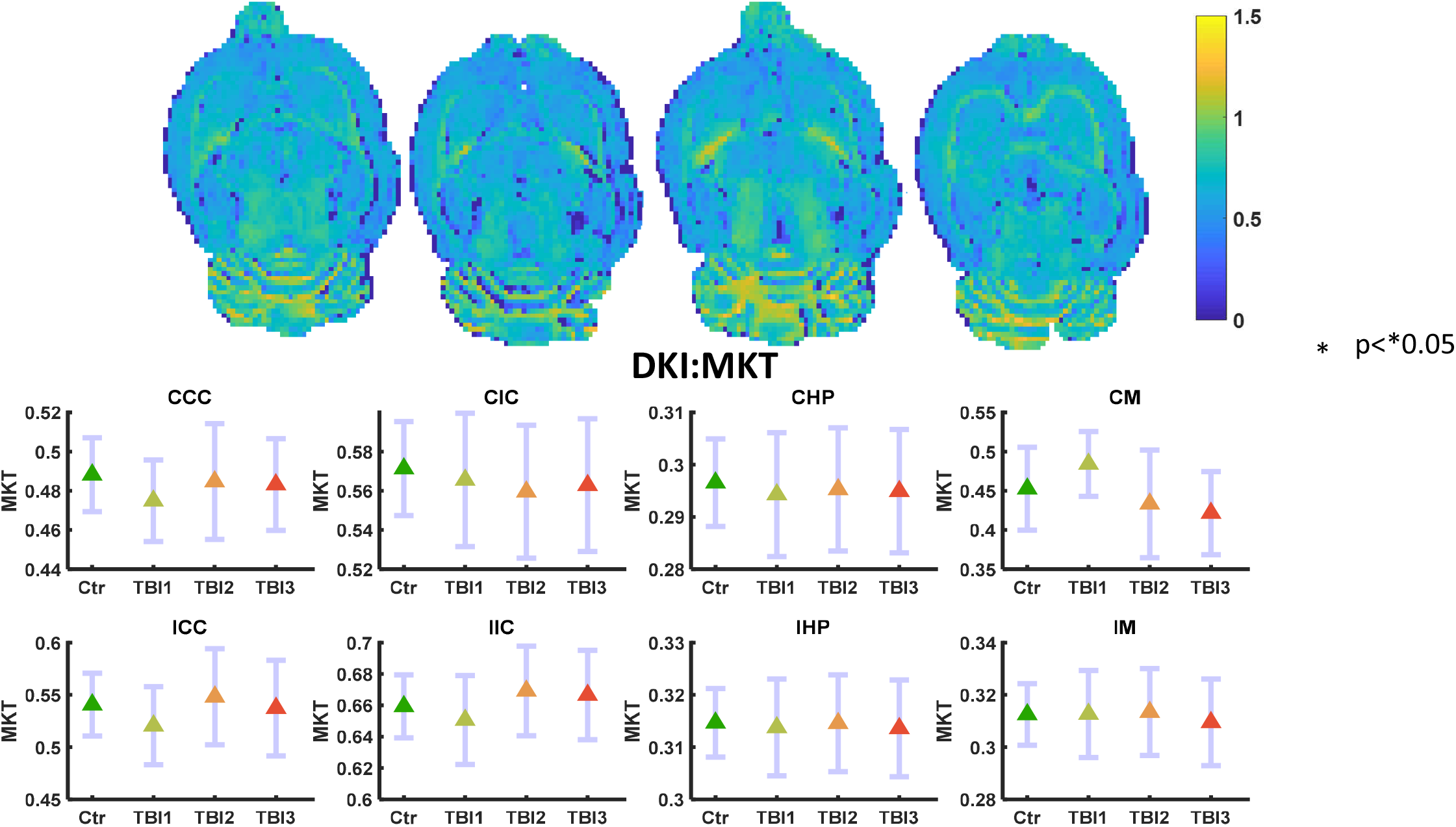
MKT maps (L-R) control, TBI1, TBI2, and TBI3. Mean (middle) and 95% confidence interval (CI) as error bar in contralateral and ipsilateral ROIs. MKT did not change significantly in any region of the brain.

#### 3.1.2 DDE Parameters (AD, RD, MD, and FA)

DDE parameters, ε, FE and MA did not show any significant alterations even before FDR correction. Higher variance (noisier maps) were observed in the higher order (*q*^4^) diffusion metrics as compared to the DTI metrics (*q*^2^) based on the DDE dataset. Among DTI parameters calculated from DDE dataset, only MD showed significant change (increase) in the ICC region (FDR corrected *p<0.05) of the brain after the third injury (Figure 7). A marked increase in DDE MD was also observed in the CCC and CHP regions (Uncorrected *p<0.05) of the brain after the third mTBI, but this did not survive FDR correction. No other ROIs showed any significant alterations at any other time points of the study (Figure 8–10).

**Figure 7:**
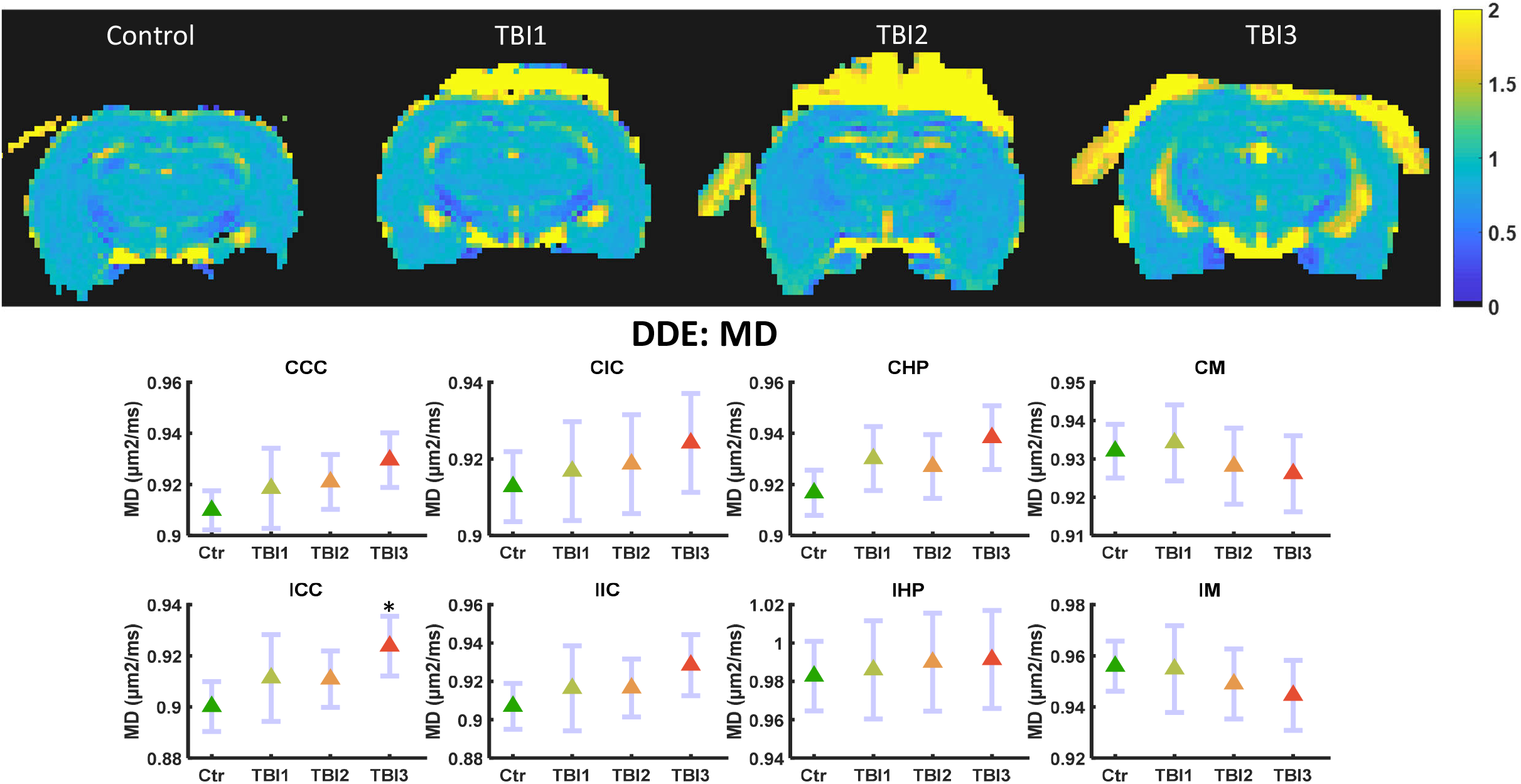
DDE data-based MD maps (L-R) control, TBI1, TBI2, and TBI3. Mean (middle) and 95% confidence interval (CI) as error bar in contralateral and ipsilateral ROIs. MD significantly increased in ICC (FDR corrected*p<0.05) region of the brain.

**Figure 8:**
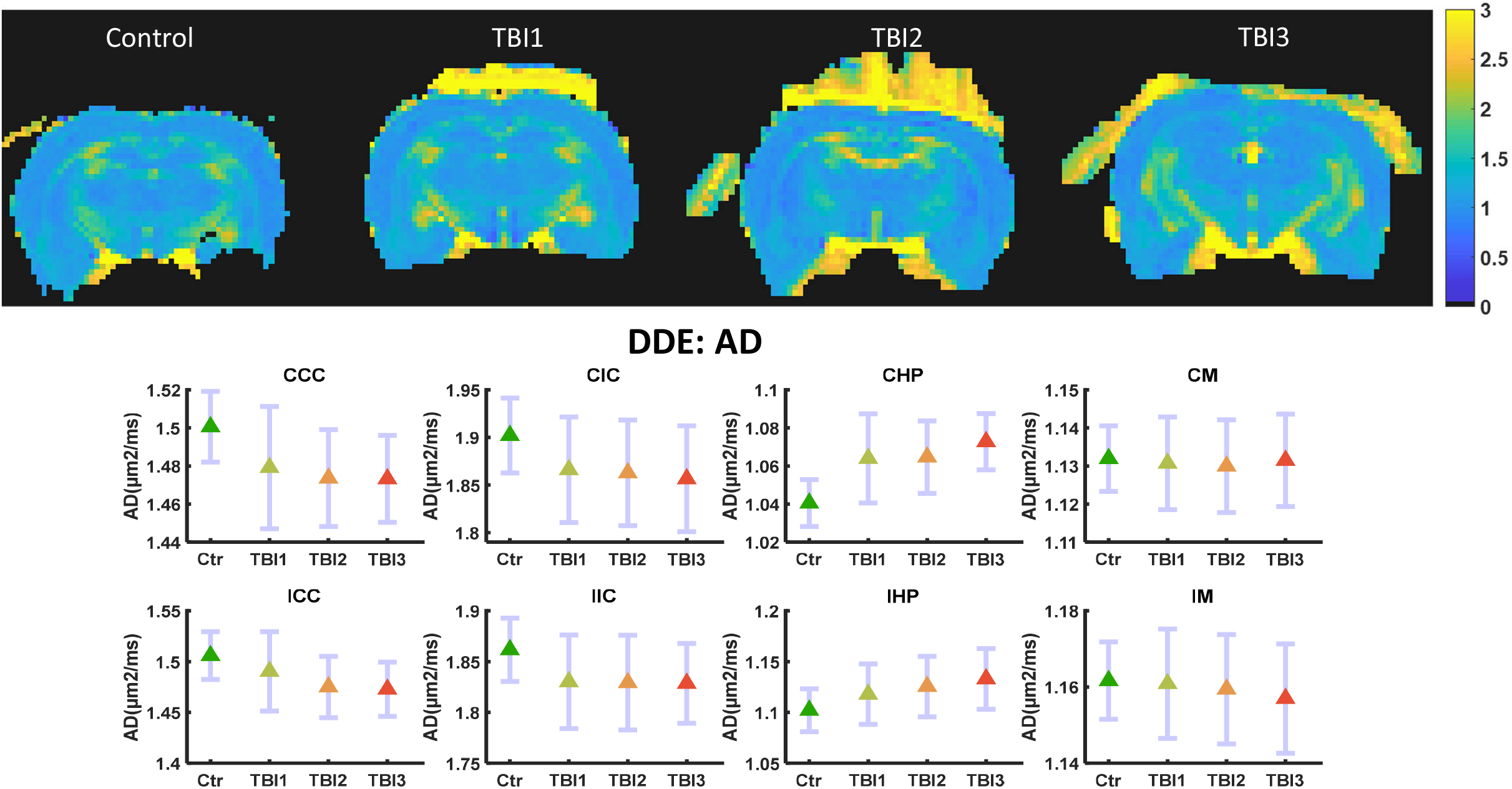
DDE data-based AD maps (L-R) control, TBI1, TBI2, and TBI3. Mean (middle) and 95% confidence interval (CI) as error bar in contralateral and ipsilateral ROIs. AD did not change significantly (FDR corrected) in any ROI of the brain.

**Figure 9:**
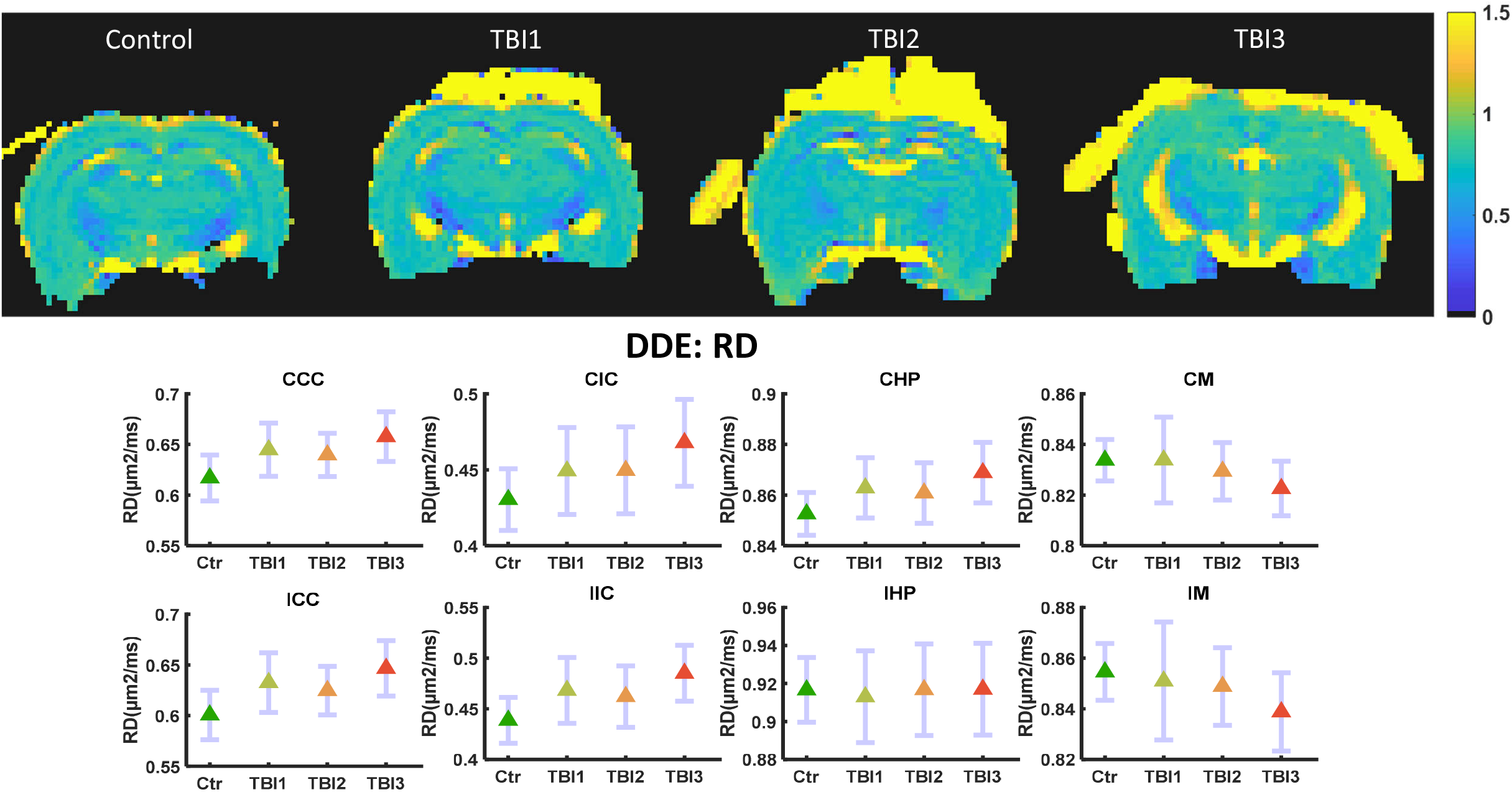
DDE data-based RD maps (L-R) control, TBI1, TBI2, and TBI3. Mean (middle) and 95% confidence interval (CI) as error bar in contralateral and ipsilateral ROIs. RD did not change significantly (FDR Corrected) in any ROI of the brain.

**Figure 10:**
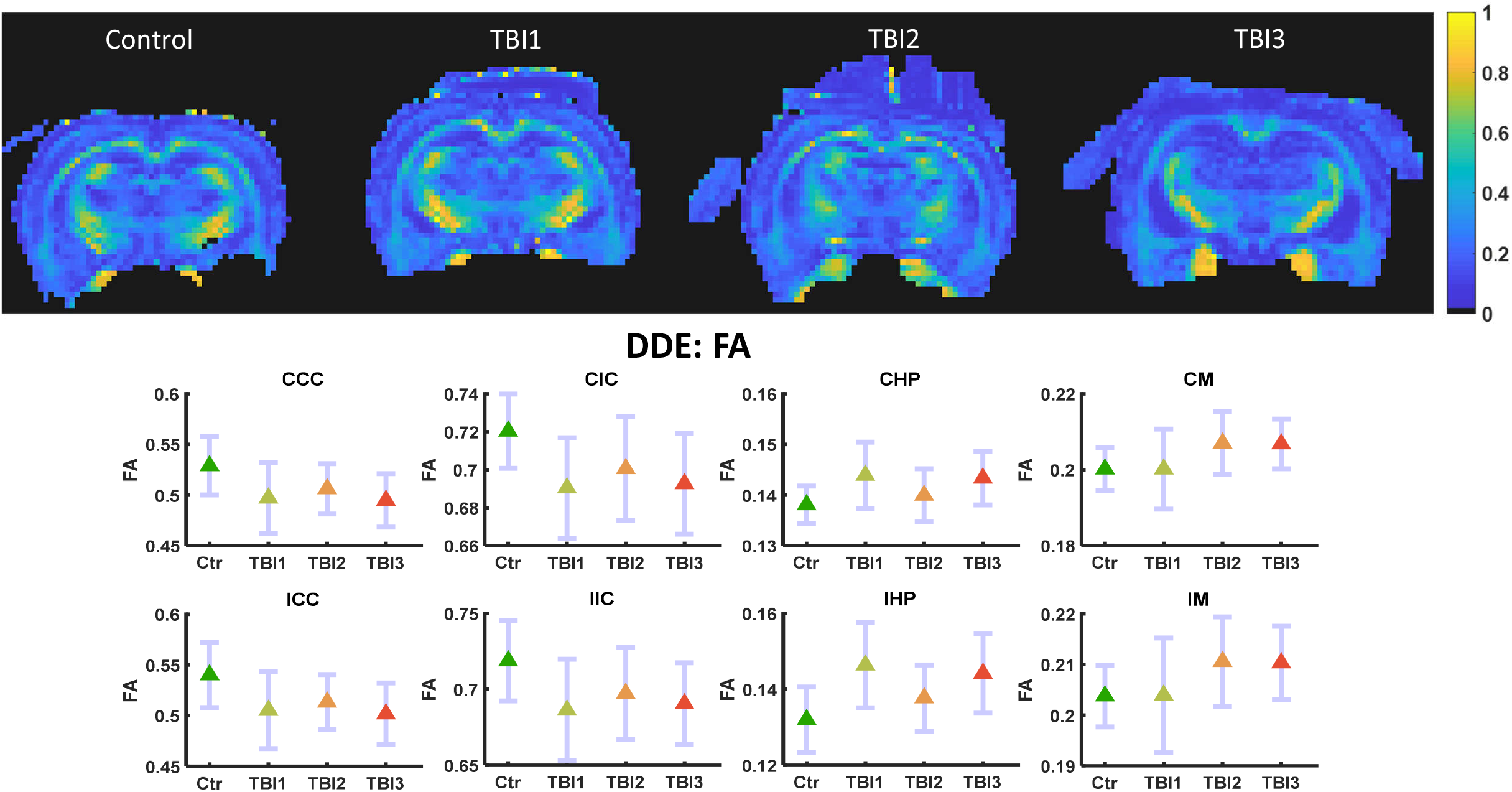
DDE data-based FA maps (L-R) control, TBI1, TBI2, and TBI3. Mean (middle) and 95% confidence interval (CI) as error bar in contralateral and ipsilateral ROIs. FA was not significantly altered (FDR corrected) in any ROI of the brain.

### 3.2 ^1^H MRS

Of the eight metabolites surviving the CRLB<20% criteria, three were found to be significantly altered after repetitive mTBI. GSH/tCr decreased markedly compared to the baseline even after the first mTBI. However, GSH/tCr was significantly decreased in comparison to baseline only after second and third mTBI (FDR corrected *p<0.05) (Figure 11). In contrast, tCho/tCr was significantly increased after second and third mTBI (FDR corrected *p<0.05) compared to baseline, and also displayed a marked increase already after the first mTBI. Ins/tCr decreased significantly after the second injury (FDR corrected *p<0.05), but not after the third mTBI in comparison to the baseline. Similarly, Tau/tCr also showed a pattern of alterations in comparison to baseline, but this was not significant. There were marked increase in Glu/tCr, GABA/tCr, and NAA+NAAG/tCr was also increased in comparison to baseline, however it did not remain significant after FDR correction (Figure 11).

**Figure 11:**
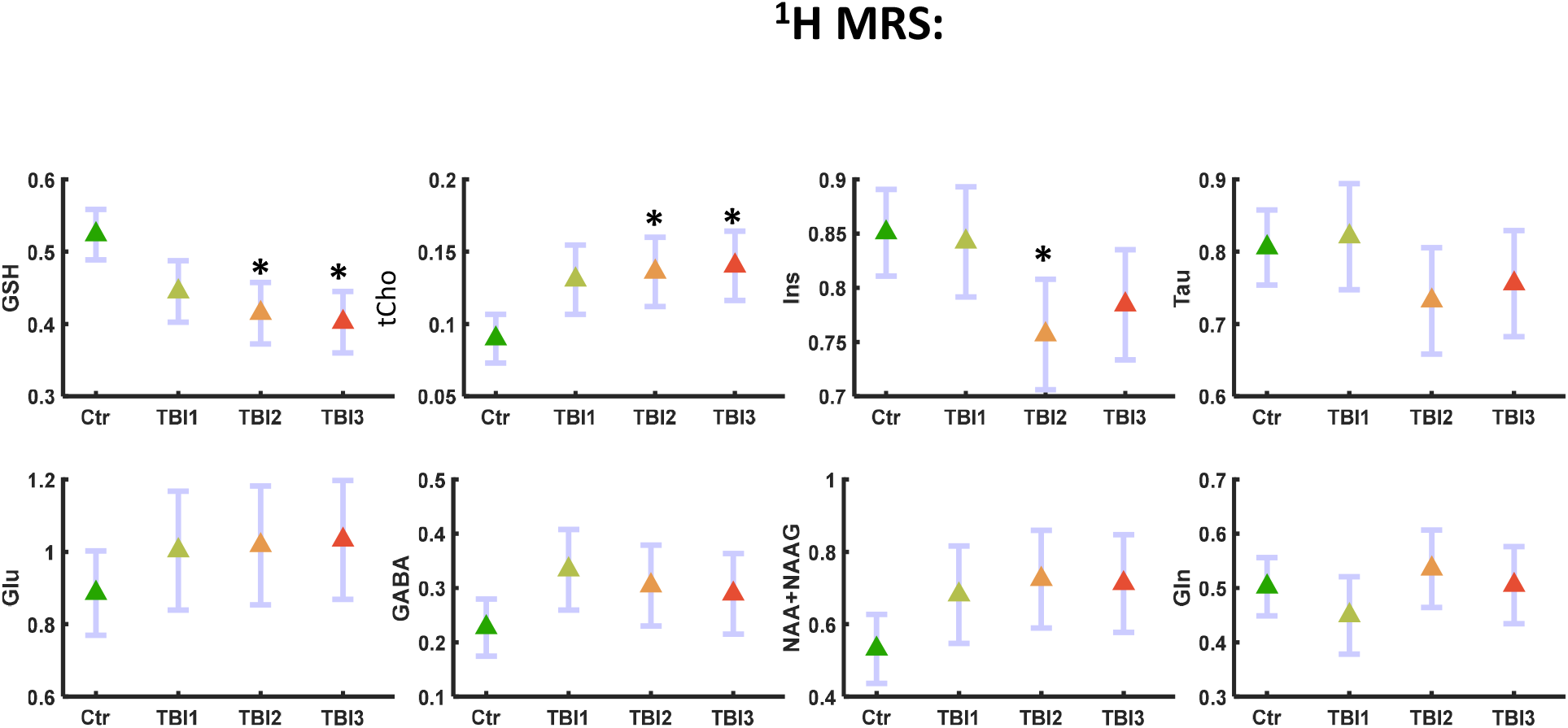
^1^H MRS based metabolites ratios with tCr (L-R) in control, TBI1, TBI2, and TBI3. Mean (middle) and 95% confidence interval (CI) as error bar in contralateral and ipsilateral ROIs. GSH significantly decreased and tCho significantly increased (FDR corrected *p<0.05) after TBI2 and TBI3. Ins significantly decreased (FDR corrected *p<0.05) only after TBI2. Glu, GABA and NAA+NAAG markedly increased in comparison to control, however, not significant.

### 3.3 Perfusion MRI (CBF)

Pseudo-continuous arterial spin labeling (pCASL) based CBF maps showed a marked decrease in the ipsilateral gray matter (IGM) and contralateral gray matter (CGM) region of the brain in comparison to the baseline maps, which did not survive FDR correction. There were no marked alterations in the CBF of WM in both ipsilateral and contralateral region of the brain, however, a marked spatial heterogeneity in the CBF was observed after second and third mTBI in all ROIs (Figure 12).

**Figure 12:**
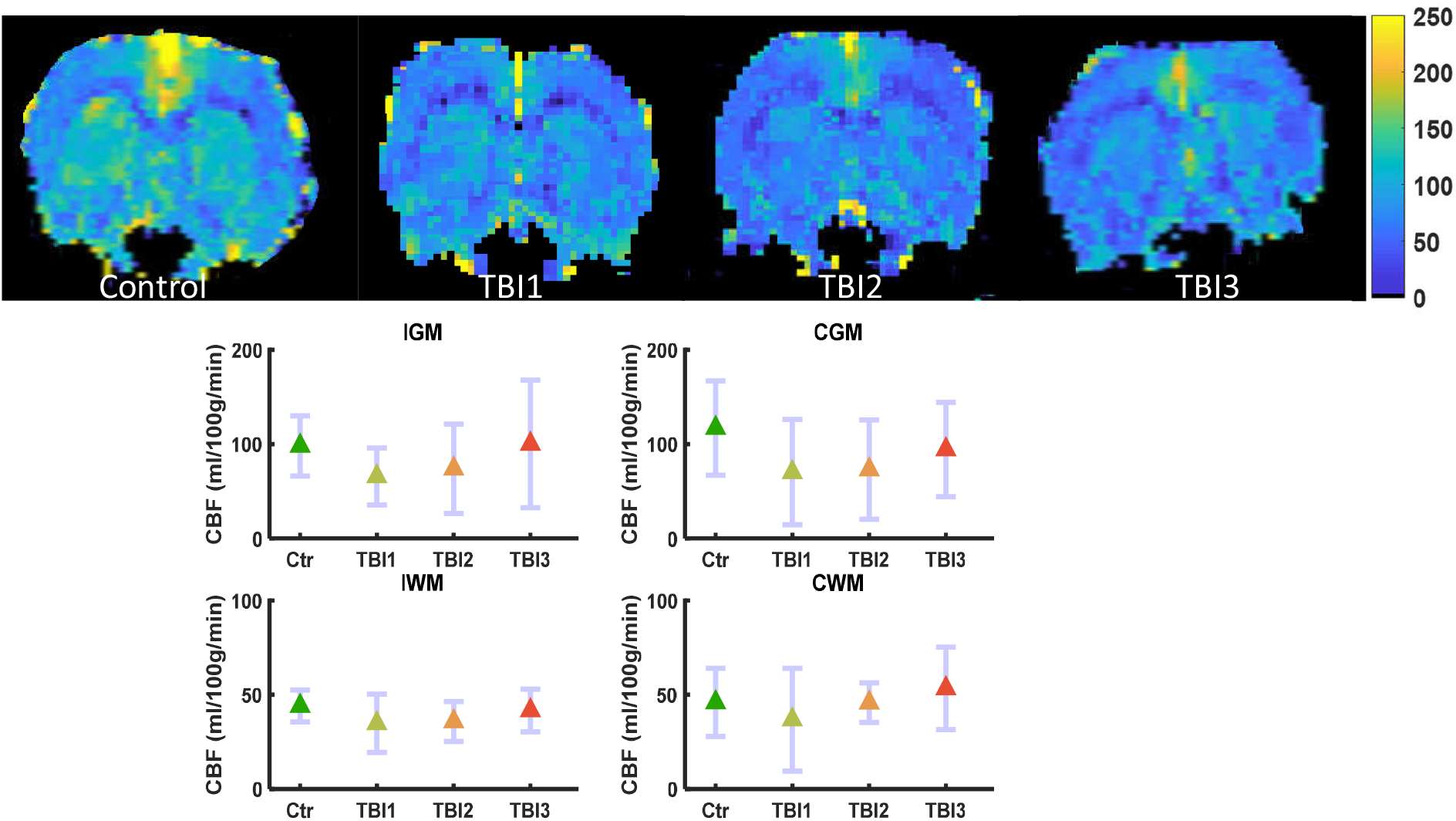
Perfusion MRI based CBF maps (L-R) control, TBI1, TBI2, and TBI3. Mean (middle) and 95% confidence interval (CI) as error bar in contralateral and ipsilateral grey and white matter ROIs. CBF was not significantly altered in any ROI of the brain, however, increased flow heterogeneity was observed after TBI2 and TBI3 in ipsilateral grey matter (IGM) region of the brain.

## 4.0 Discussion

In the last few decades clinical, and preclinical research has identified potential neuroimaging biomarkers for repetitive/ mTBI [39, 40], but until now an objective diagnostic test is not available due to immense heterogeneity of symptoms and/or inability to probe subtle metabolic, microstructural and pathophysiological alterations with traditional neuroimaging techniques. Early detection to probe metabolic, microstructural and pathophysiological alterations after repetitive/mTBI cases are crucial for treatment intervention and management.

### 4.1 DKI based findings

Among DKI based metrics, only *W*_T_ was found to have a significant decrease in IHP region: A mild decrease after first and second mTBI, and significant only after the third mTBI at day 7, which may suggest progressive demyelination after repetitive/mTBI. Similar to the present finding, a cuprizone treated rat demyelinating model also showed a significant decrease in radial kurtosis [56]. A significant decrease in *W*_T_ may also be due to the substantial spine loss, dendritic and axonal beading or fragmentation after mTBI [11]. However, fragmentation or axotomy would likely be accompanied by marked alterations in AD and in *W*_L_, which was found neither in the gray matter nor the white matter regions of interest. However, repeated mTBI may induce an energy deficient, edematous, ischemic and/or metabolically compromised state, leading to persistent dendritic beading and loss of dendritic spines. These factors may lead to a significant decrease in *W*_T_ in the IHP, but not in CHP region after repetitive mTBI at day7. This sensitivity to injury of the hippocampus region is well-know and has been demonstrated by even mild stressors [57]. A simulation and sensitivity analysis of acute nervous-system injury has investigated the effect of edema and beading on dMRI data for better understanding and interpretation [12], and supports the speculation that significant decrease in *W*_T_ may serve as an indicator of demyelination, dendritic beading, edema, or a mixture of such pathologies after repetitive mTBI.

Another repetitive mTBI study [58] using DKI also reported no significant alteration in the white matter region up to day 6 after repetitive mTBI in comparison to the sham control, corroborating the present DKI findings in white matter regions. However, [58] reported significantly decreased FA and AD and a significantly increased AK on day 6 in the cortex with significant astrogliosis, while we did not observe any significant alteration in SDE-DTI parameters in any ROIs. A significant correlation of mean kurtosis with astrogliosis in an mTBI rat model [59] further supports the enhanced sensitivity of DKI towards subtle microstructural alterations in comparison to the DTI. The DKI findings of the present study are also unique in that unlike other repetitive/mTBI studies, anatomical scans did not show any apparent tissue injury in the ipsilateral cortical region of the brain directly beneath the impact site of the brain. This is similar to human mTBI scans which are often reported normal under conventional neuroimaging [4, 8].

### 4.2 DDE based findings

The three DDE metrics considered in this study did not show any significant alterations in any region of the brain at all three-time points. This might be due to the higher order diffusion effect, which generally appeared noisier (higher variance in the fourth order in q DDE metrics than the DDE based DTI metrics). This was also reported in [26]. However, the DDE based DTI MD, which distinguishes itself from its SDE counterpart mainly by a shorter diffusion time and a longer echo time, showed a significant increase in ICC region of the brain. Increase in the MD and decrease in the FA (mild decrease in the present study) could be attributed to reduced cellular density in the gray matter region and fiber density in the white matter region. Additionally, the present study also found a marked decrease in AD in the white matter ROIs and a slight increase in subcortical gray matter ROIs, also corroborating previous mTBI studies [11, 12, 41, 60]. A significant increase in MD in ICC and CCC region was also reported in a rat model of repetitive fluid percussion mTBI and found to normalize after treatment [61]. These findings suggest that the corpus callosum is a sensitive region for repetitive/ mTBI and such consistent findings may be helpful in the diagnosis of repetitive mTBI. However, it should be kept in mind that most of the previous mTBI studies employed an mTBI paradigm exhibiting macroscopic neural damage, while the present study did not show any conspicuous neural tissue injury.

### 4.3 ^1^H MRS

Among eight metabolites analyzed from ^1^H MRS spectrum (<20 % CRLB criteria), three metabolites (GSH, Cho, and Ins) showed significant alterations, but only after repetitive mTBI. Thus, no significant alteration was observed after the initial mTBI. GSH was significantly decreased after the second and third mTBI in comparison to the baseline. A significant decrease in GSH may be due to the increase in reactive oxygen/ nitrogen species (ROS/NOS) after repetitive mTBI. Depletion of GSH is also related to an increase in cyclooxygenase-2 activity, which promotes neuroinflammation and lipid peroxidation. Lipid peroxidation may induce cellular vulnerability of increased susceptibility of repetitive injury [62].

A progressive increase in Cho after repetitive mTBI in the present study also suggests lipid peroxidation induced membrane breakdown and/or repair. A progressive increase in tCho compounds at the injury site may be due to the release of acetylcholinesterase, which converts acetylcholine to choline and acetate which has been reported in previous mTBI studies [63, 64]. Higher tCho concentration in comparison to baseline may also suggest heightened cell membrane turnover, as reported in demyelination, remyelination, inflammation, and gliosis [65]. Previous studies have also reported an increase in Cho/Cr level after repetitive/mTBI studies in human [66]. However, a mild TBI study in humans has reported low Cho/Cr at the sub-acute stage of mTBI [66]. Nonetheless, a biphasic response of neurometabolites at the acute, sub-acute, or chronic phase has been reported previously [66]. Therefore caution is needed when comparing metabolic level estimates to the phase of repetitive/mTBI. Our findings of increased choline, Glu– Gln levels (which may indicate diffuse axonal injury and excitotoxicity, respectively), and cerebral metabolites associated with neuroinflammatory changes, such as mI are also corroborated by previous mTBI studies using ^1^H MRS [67].

### 4.4 Perfusion MRI (CBF)

The perfusion MRI based CBF parameter did not show any significant alterations in any group and in any ROIs of the brain before FDR correction. Nevertheless, we note that a marked decrease in CBF after mTBI was observed in the cortical impact region (IGM). After acute mTBI, the impact area may become hypermetabolic, but at the same time, there is a decrease in CBF [16, 68]. This mismatch of supply and demand leads to an energy crisis at the cellular level, resulting in additional death of neural tissue in the ischemic penumbra. However, at the acute phase of repetitive mTBI, we observed increased heterogeneity in the CBF map (within the IGM ROI), which may suggest increased capillary transit time heterogeneity (CTH) [69]. Such an increased CTH can lead to IGM-tissue hypoxia via shunting of oxygenated blood through the capillary bed, even in the absence of significant CBF reduction as observed in the present study. Although reductions in CBF, paradoxically, improve brain oxygenation if CTH is critically elevated [37]. The contributing factors for increased CTH are oxidative stress, vasogenic edema, leukocytosis, astrocytic endfeet swelling to name a few. Oxidative stress-induced free radicals affect capillary contraction function of pericytes and around 40% of capillary pericytes leave their pericapillary location within the first hour of the insult [34, 37]. Pericyte depletion also causes white matter dysfunction [70], potentially reflected in the significantly decreased MD in ICC in the present study. The metabolically compromised impact site presumably also shows higher lactate levels after repetitive mTBI, but lactate was not included in our analysis due to high CRLB. Increase in lactate concentrations over 5 mmol/kg were found to induce cerebral hyperemia and this may also contribute to increased CTH after repetitive mTBI [71],. Therefore, diffusion, perfusion, and ^1^H MRS parameters are complementary and interdependent and support the usefulness of multiparametric MRI.

## 5.0 Limitations

The present study reveals important MRI findings of repetitive mTBI, which can help in the diagnosis and prognosis of repetitive mTBI. However, the study has some limitations such as the absence of immunohistochemistry due to the absence of a separate control group. The higher order DDE metrics did not show any marked alterations, maybe due to a low SNR. However, higher DDE SNR is certainly possible through averaging and so DDE should not be ruled out in future studies. Indeed, DDE metrics were suggested to have improved sensitivity for age-related microstructural changes in a human population [24]. The DKI dataset did not shown a significant alteration in DTI parameters may be due to limited diffusion directions or due to the very mild nature of mTBI protocol employed here. The ^1^H MRS has been acquired only from the ipsilateral cerebral cortex, however, multivoxel spectroscopy can reveal metabolic alterations in other ROIs also, which may provide complementary information related to the tissue microstructural alterations in the IHP. Perfusion MRI was analyzed only in a single slice covering the injury site, and can therefore not address potential changes elsewhere. Future studies should employ perfusion methods that could provide CBF information of the whole brain so that CBF values of the targeted ROIs can be extracted and evaluated in comparison to the control. Only five animals were employed in this longitudinal study to explore microstructural, metabolic and, pathophysiologic alterations, however, more experimental animals would certainly be advantageous for a multiparametric study.

## 6.0 Conclusion

The present study employs a multiparametric MRI approach which reveals significant microstructural, metabolic and marked CBF alterations after repetitive mTBI. A longitudinal evaluation of such changes after mTBI also supports the vulnerability of the brain towards repetitive impacts. Particularly, significant microstructural alterations in IHP and ICC after repetitive mTBI supports the sensitivity of these brain regions towards mTBI. ^1^H MRS based metabolic measurements in the ipsilateral cerebral cortex also revealed significant alteration in antioxidant level (GSH) and cellular turnover marker (tCho) at the impact site after repetitive mTBI, suggesting increased oxidative stress due to repetitive mTBI. The marked reduction in CBF after the first mTBI and subsequently increased confidence interval of the CBF also indicate aggravated pathophysiology. Each MRI parameter independently indicated some aspect of the pathological characteristics of repetitive mTBI. Therefore, a multiparametric MRI approach provides complementary information, which can be used as potential biomarkers in diagnosis or prognosis of normal appearing repetitive mTBI cases.

## ACKNOWLEDGEMENT

Danish Ministry of Science, Technology and Innovation’s University Investment Grant (MIND Lab, Grant no. 0601–01354B), and NIH 1R01EB012874-01. Danish Research Council’s Infrastructure program, the Velux Foundations, and the Department of Clinical Medicine, AU.

