## Supplementary material for "Longitudinal, Multiparametric MRI Assessment of repetitive mild TBI in rats": Supplemetary_Material

**Supplementary Video 1**

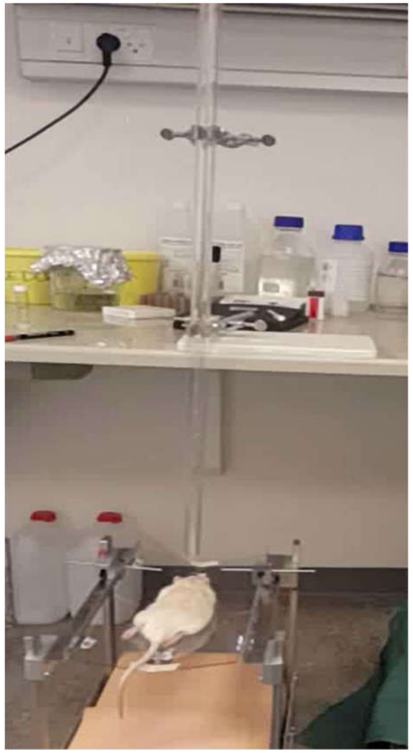

**Supplementary figure 1(A)**

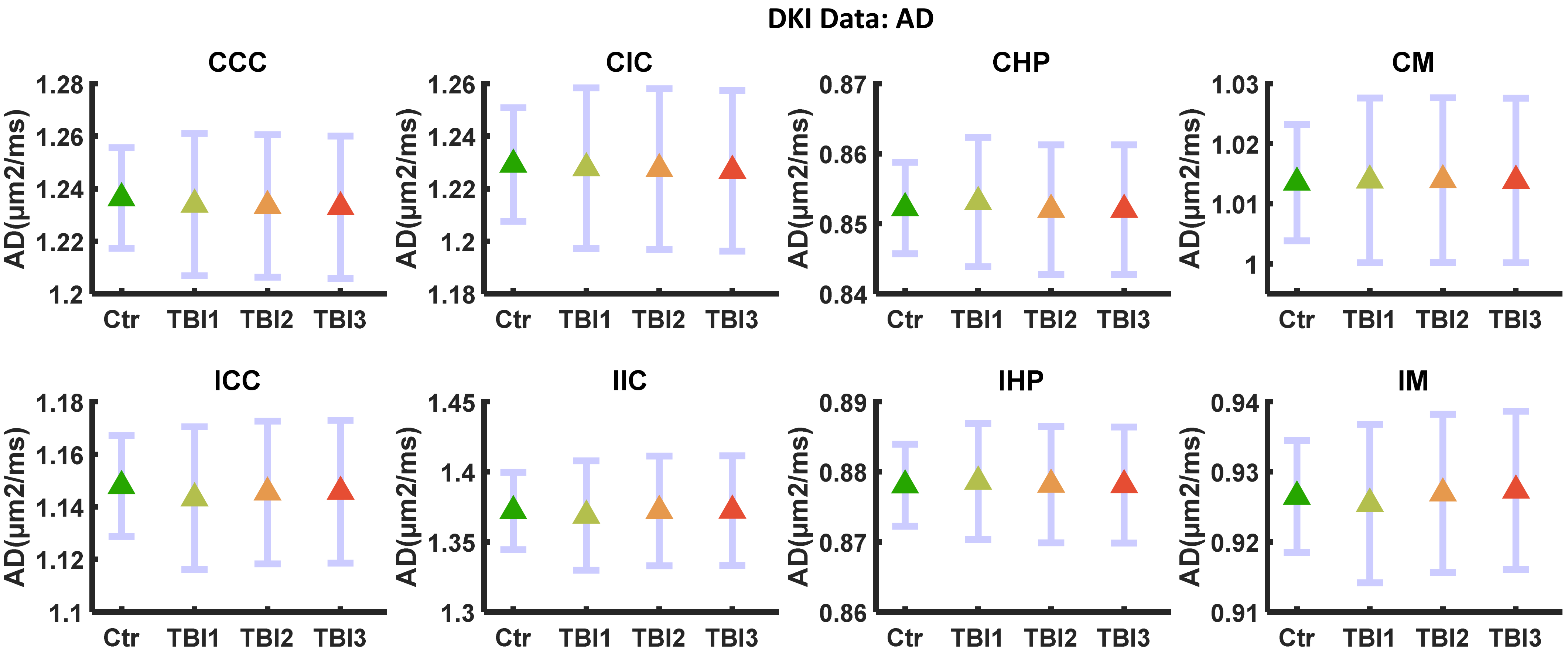

**Supplementary figure 1(B)**

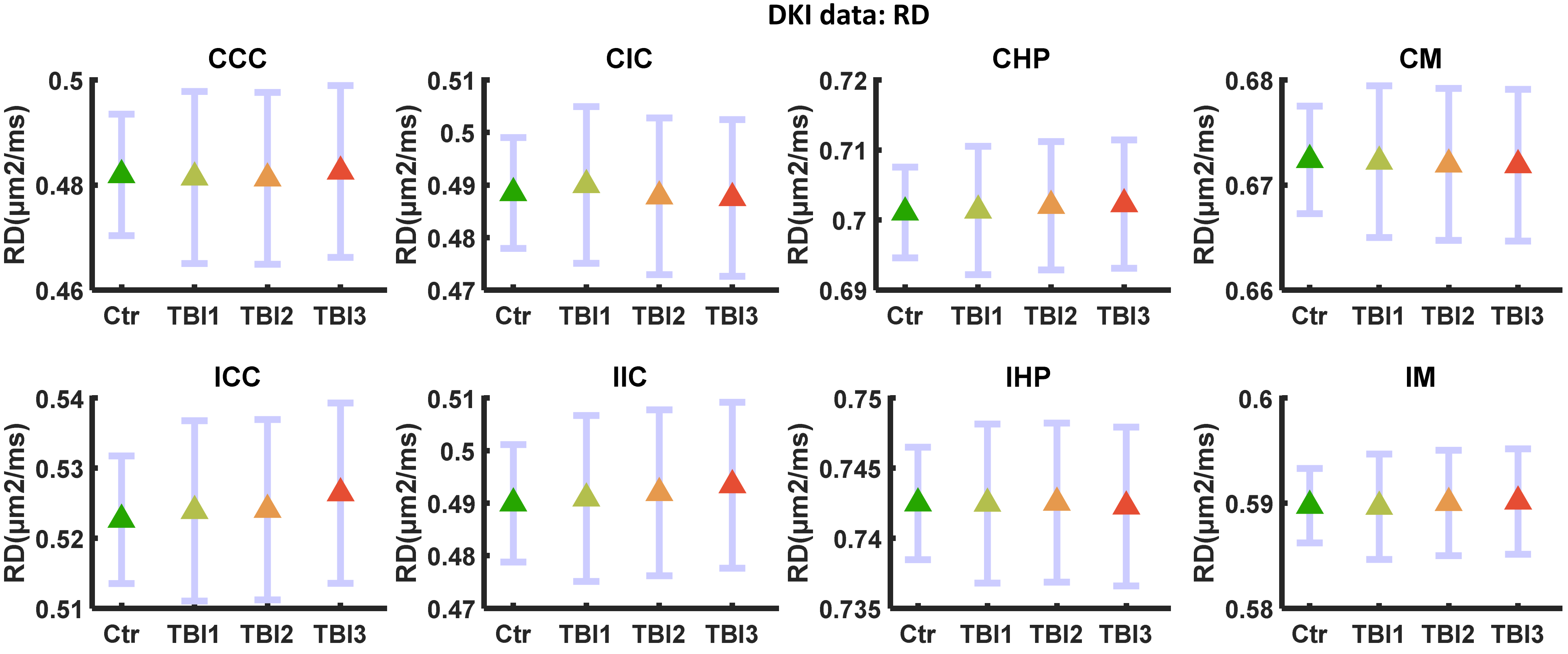

Supplementary figure 1(C)

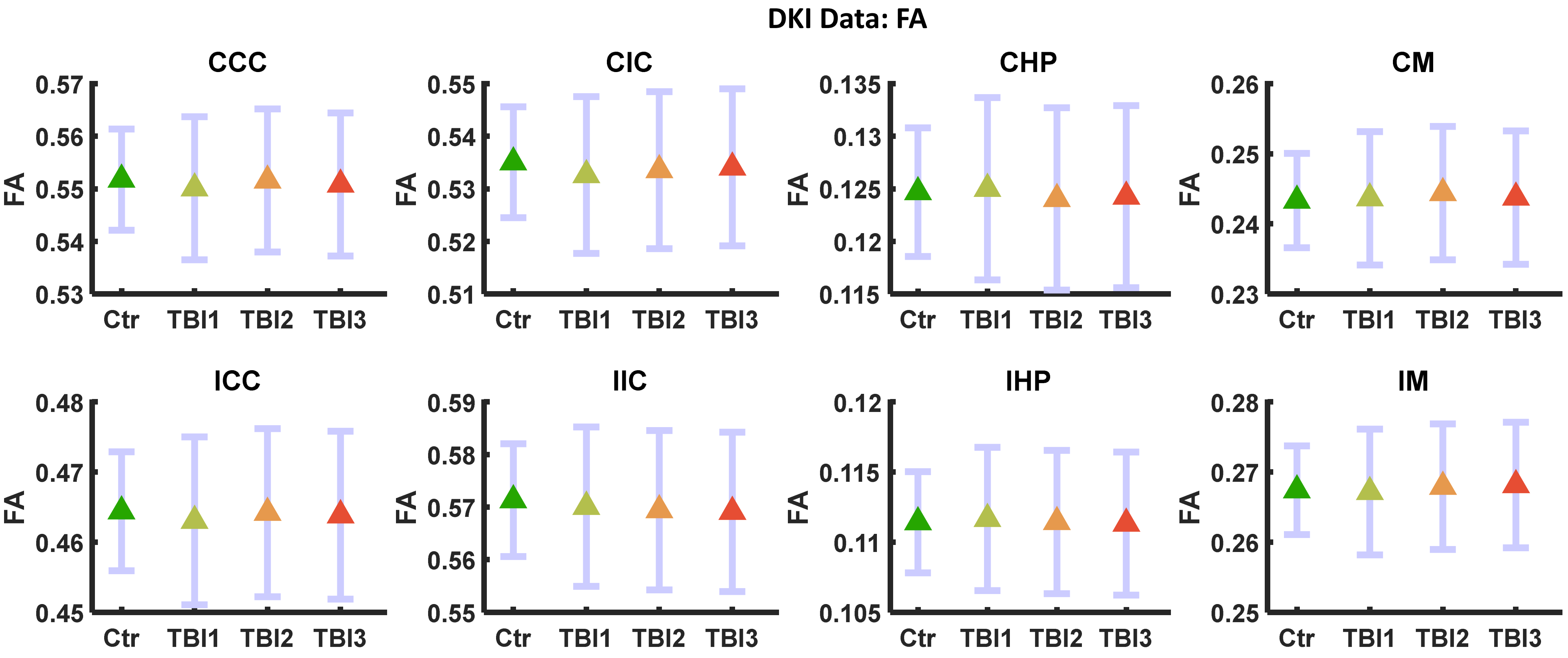

**Supplementary figure 1(D)**

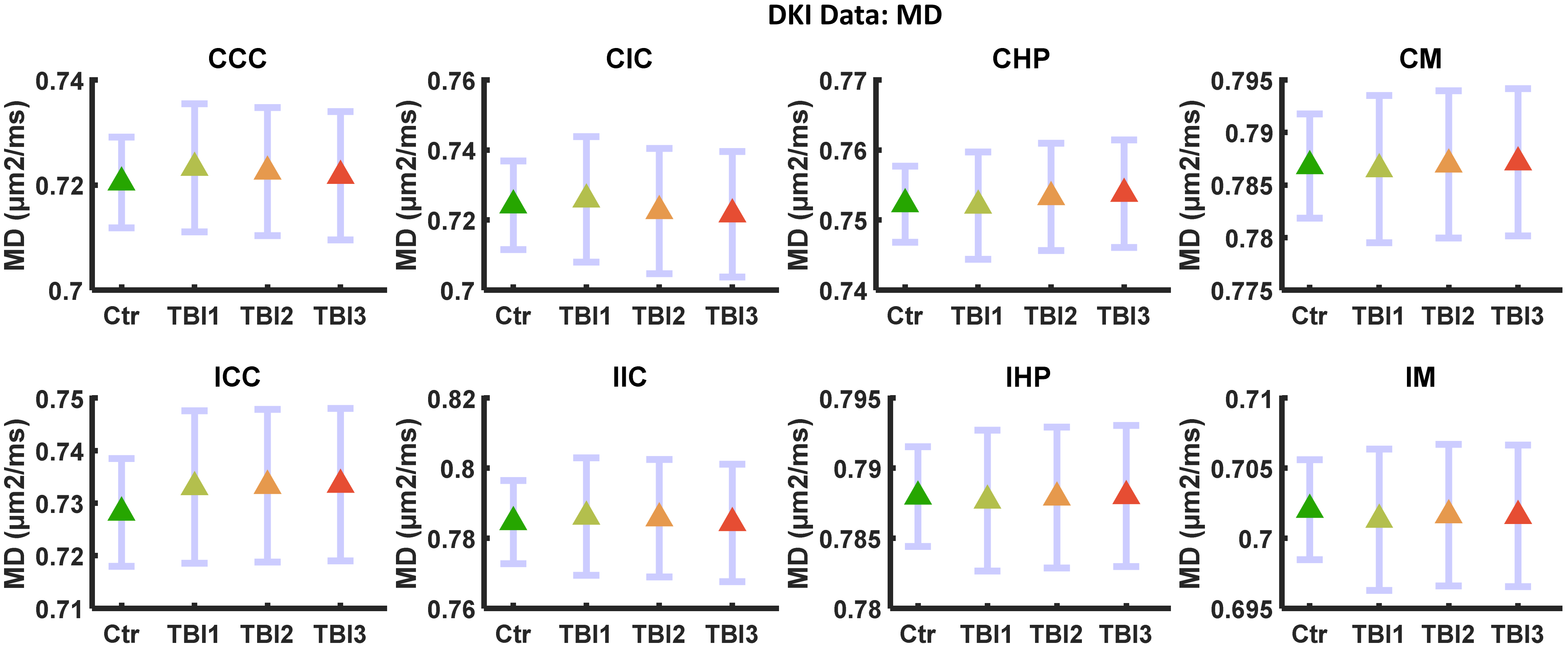

### Supplementary figure 2(A)

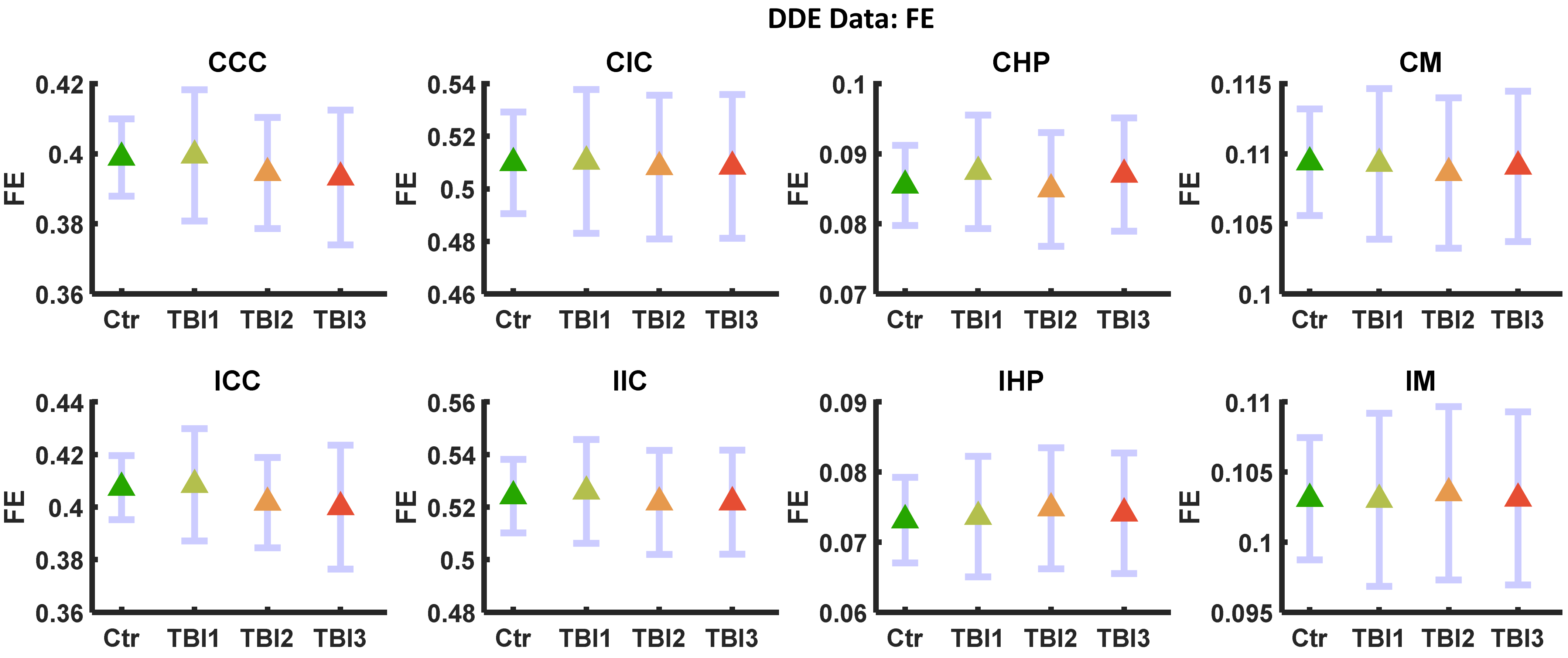

### Supplementary figure 2(B)

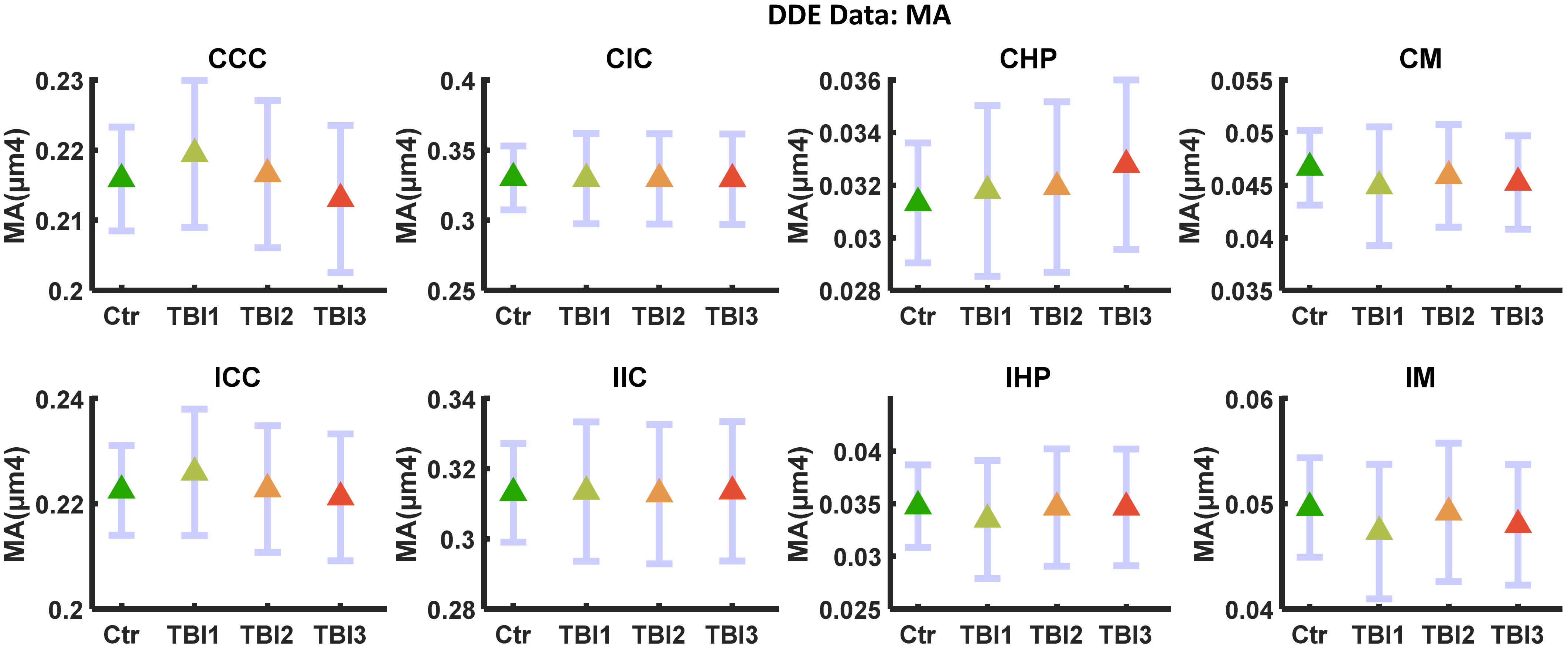

**Supplementary figure 2(C)**

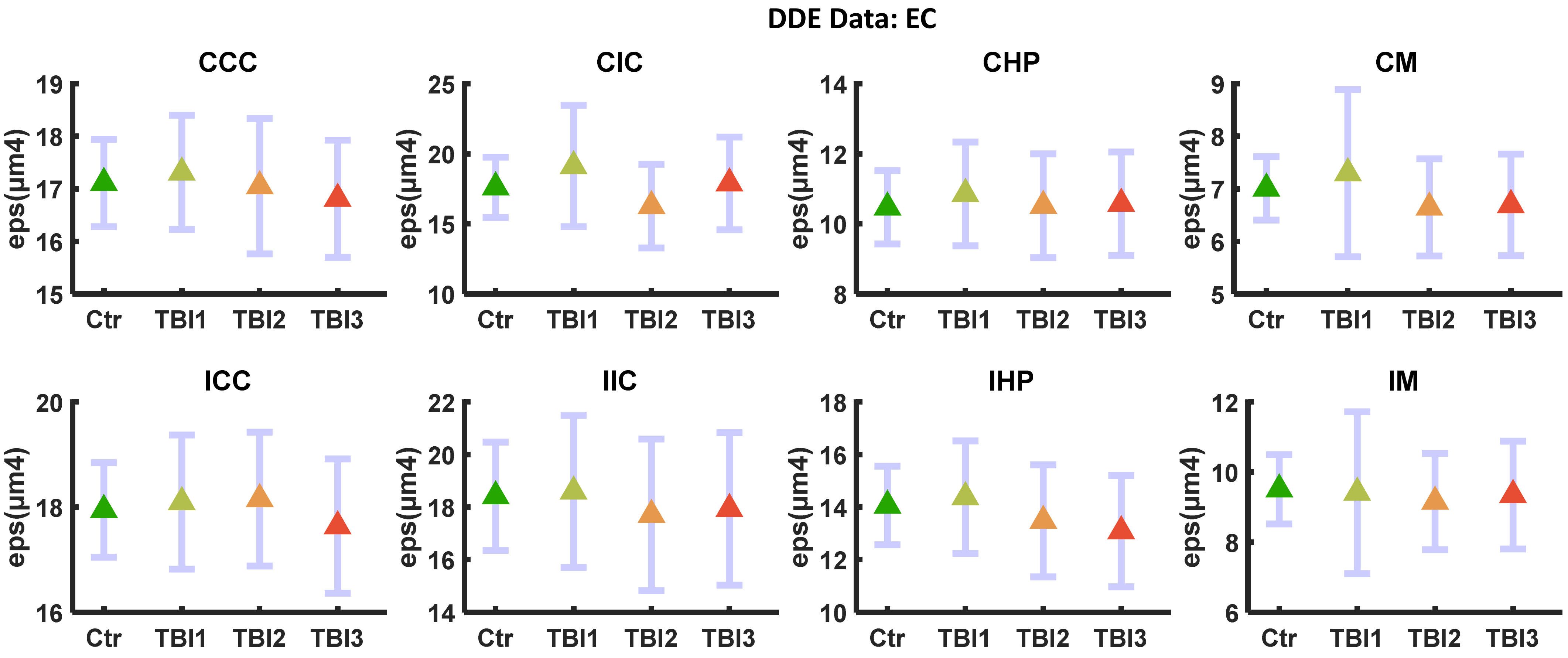

**Supplementary figure 3**

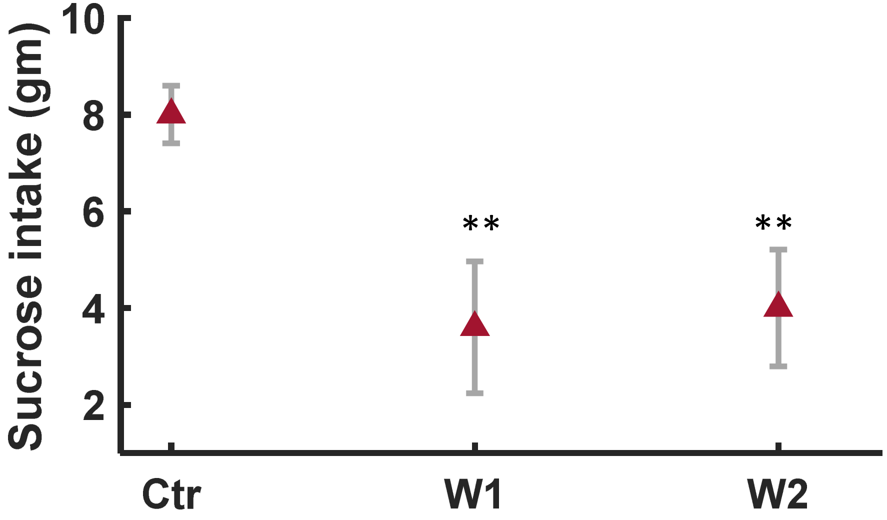


**p<0.01
